# Engineering human spinal microphysiological systems to model opioid-induced tolerance

**DOI:** 10.1101/2022.10.03.510710

**Authors:** Hongwei Cai, Zheng Ao, Chunhui Tian, Zhuhao Wu, Connor Kaurich, Zi Chen, Mingxia Gu, Andrea G. Hohmann, Ken Mackie, Feng Guo

## Abstract

Opioids are commonly used for treating chronic pain. However, with continued use, they may induce tolerance and/or hyperalgesia, which limits therapeutic efficacy. The human mechanisms of opioid-induced hyperalgesia are significantly understudied, in part, because current models cannot fully recapitulate human pathology. Here, we engineered novel human spinal microphysiological systems (MPSs) integrated with plug-and-play neural activity sensing for modeling human nociception and opioid-induced tolerance. Each spinal MPS consists of a flattened human spinal cord organoid derived from human stem cells and a 3D printed organoid holder device for plug-and-play neural activity measurement. We found that the flattened organoid design of MPSs not only reduces hypoxia and necrosis in the organoids, but also promotes their neuron maturation, neural activity, and functional development. We further demonstrated that prolonged opioid exposure resulted in neurochemical correlates of opioid tolerance and hyperalgesia, as measured by altered neural activity, reduced densities of glutamate transporter levels and downregulation of μ-opioid receptor expression of human spinal MPSs. The MPSs are scalable, cost-effective, easy-to-use, and compatible with commonly-used well-plates, thus allowing plug-and-play measurements of neural activity. We believe the MPSs hold a promising translational potential for studying human pain etiology, screening new treatments, and validating novel therapeutics for human pain medicine.

## 1. Introduction

More than 20% of the general population suffers from chronic pain.^1, 2^ Due to the increasing frequency of chronic pain and its heavy burden on patients and society, there is increasing interest in studying pain mechanisms, treatment, and management.^3^ Opioids are one of the most commonly prescribed medications for pain.^4, 5^ However, the administration of opioids causes undesired side effects, including opioid-induced tolerance and hyperalgesia^6, 7^ which limits their efficacy to relieve pain over time. The first clinical report of opioid-induced tolerance can be traced back to the 19th century.^5^ More recently, human subjects and animal models have been used to study the complicated pathology of opioid tolerance and hyperalgesia^6, 7^. Several human trials have been conducted to examine the existence and potential clinical significance of opioid-induced tolerance during chronic opioid therapy.^8-11^ This approach is limited by experimental accessibility as well as moral and health concerns. Animal models have been extensively used for studying the mechanism(s) and pathology underlying opioid-induced tolerance.^12^ For example, animal spinal cord slices were used to discover neuronal activity and receptor changes related to opioid-induced tolerance.^13-15^ However, a challenge remains in translating discoveries from animal models to human treatment due to the different biology between animal models and humans,^16^ and concerns about the translational relevance of preclinical pain models. Thus, there is an urgent and unmet need to develop models for understanding opioid-induced tolerance in humans.

Recent advances in stem cell and organoid technologies show promise for modeling human pain and opioid-induced tolerance.^17^ Human organoids are organ-like 3D in vitro cultures derived from human stem cells, recapitulating key neuroanatomical and neurochemical features observed in vivo. Several pioneering studies have been reported to develop human spinal cord organoids and sensory neurons. For example, human sensory neurons^18^ and sensory ganglion organiods^19^ have been differentiated from induced pluripotent stem cells (iPSCs) for pain research. Human ventral spinal cord organoids have been developed to study motor neuron development and activity, and cortex-spinal-muscle assemblies were generated to study motor neuron innervation of muscles in vitro.^20^ Moreover, protocols to generate spinal cord organoids with dorsal interneurons and sensory neurons have been developed for modeling spinal cord disease and neural activity.^21, 22^ However, limitations of current human spinal cord organoids hinder their applications for studying human neural activity and pathology of opioid-induced tolerance including the fact that (1) current human spinal cord organoids are highly heterogeneous; (2) current human spinal cord organoids suffer from hypoxia and necrosis, significantly affecting neural activity; and (3) it is hard to obtain a stable and reproducible neural electrical measurement of human spinal cord organoids due to the organoid-electrode contact issue, and the hypoxia- and necrosis-induced neuron death during the measurement.

To address the above technical barriers and to grow better organoids, microphysiological systems and organoids-on-a-chip systems have been developed. This technology permits both the regulation and monitoring of organoid development through the integration of microfluidics and various bioengineering approaches.^23-25^ For example, efforts have been made to improve the cultural conditions of developing organoids. Pillar-based perfusion microfluidic channels have been fabricated for the on-chip formation of brain and liver organoids derived from human-induced pluripotent stem cells (iPSCs).^26-28^ Recently, a very innovative lung-on-a-chip design has been reported for mimicking lung function by microfabricating two microfluidic channels separated by a stretchable porous PDMS membrane sandwiched in-between the channels.^29-31^ The lung-on-a-chip devices and membrane-based microfabricated devices have been adopted to improve the perfusion of oxygen and nutrients and generate various types of organoids, including brain, pancreas, liver, and glomerulus organoids.^32-42^ Recently, we employed 3D printed scaffolds to generate tubular human brain organoids for reducing hypoxia/necrosis and improving the standardization of organoids.^43^ In addition to growing better organoid cultures, several attempts have been made to enable monitoring of development as well as the response to drug treatment. For example, long-term four-dimensional light-sheet microscopy has been established for recording the spatial lineage of developing cerebral organoids.^44^ Moreover, real-time live-cell imaging has been incorporated into an automated microfluidic platform for high throughput drug screening by testing the relative apoptosis and viability of tumor organoids under different drug treatments.^45^ So far, the above-mentioned pioneering works have improved the development or/and characterization of human organoids. However, there is a tremendous need to develop better models that recapitulate both human neural activity and the pathology of opioid-induced tolerance, because of technical barriers to simultaneously maintaining high-quality human spinal cord organoids and stably measuring neural activity in response to opioids.

Herein, we report novel human spinal microphysiological systems (MPSs) integrated with plug-and-play neural activity sensing to model human biology underlying opioid-induced tolerance and hyperalgesia. To avoid hypoxia/necrosis within conventional organoids, we innovated a flattened organoid design of MPSs to enhance nutrient and oxygen perfusion. We found that the uniquely flattened organoid design not only supports neuron maturation/activity but also provides better contact between organoids and multi-electrode array (MEA) electrodes, improving the stability and reproducibility of neural activity measurement. After the integration of 3D printed organoid holder devices, the MPSs could be used for plug-and-play neural activity measurement, and are compatible with commonly-used multiwell-plates and MEA systems. As a proof-of-concept application, the human spinal MPSs were used to model neural activity and the biology of nociception and hyperalgesia induced by opioids. We demonstrated that the neural activity of human spinal MPSs was rapidly suppressed by acute opioid (DAMGO) administration and enhanced by chronic opioid exposure. Long-term DAMGO treatment of the human spinal MPSs also reduced both glutamate transporter and μ-opioid receptor levels. Thus, we believe the human spinal MPSs may hold promising potential for broad applications in basic and translational pain research.

## 2. Results and Discussion

### Engineering human spinal microphysiological system to model pain relief and opioid-induced hyperalgesia

Opioids are effective analgesics, however, chronic opioid exposure is problematic for pain management for many reasons, including tolerance and hyperalgesia. To study the latter two processes, we developed novel human spinal microphysiological systems (MPSs) integrated with plug-and-play neural activity sensing. Each human spinal MPS consists of a flattened human spinal cord organoid derived from human stem cells and a 3D printed organoid holder device for plug-and-play neural activity measurement (**Figure 1a**). The 3D printed organoid holder device (**Figure 1b**) is made by integrating a polycarbonate membrane onto a 3D printed hollow ring (outer diameter of 15 mm, inner diameter of 11 mm, and height of 5 mm), and can be directly inserted into commonly-used cell culture well-plates or multi-electrode array (MEA) systems (**Figure S1**). Since nutrients and oxygen can only penetrate about 400 μm from the surface of organoid tissue, conventional spherical organoids with a diameter up to several millimeters frequently develop central hypoxia and necrosis, detrimentally affecting neuronal function and activity. By using 3D-printed organoid holder devices, we confined the growth of human spinal cord organoids within a space (height = 500 μm) to generate flattened human spinal cord organoids.. The 500 μm organoid thickness was smaller than the double side penetration depth (∼ 800 μm), allowing sufficient nutrients and oxygen supplement. And such thickness could also support organoid structure (VZ/SVZ) development.^46^ We then used established human spinal MPSs to investigate their functions as well as cellular processes involved in hyperalgesia and tolerance induced by opioids (**Figure 1c**).

**Figure 1.**
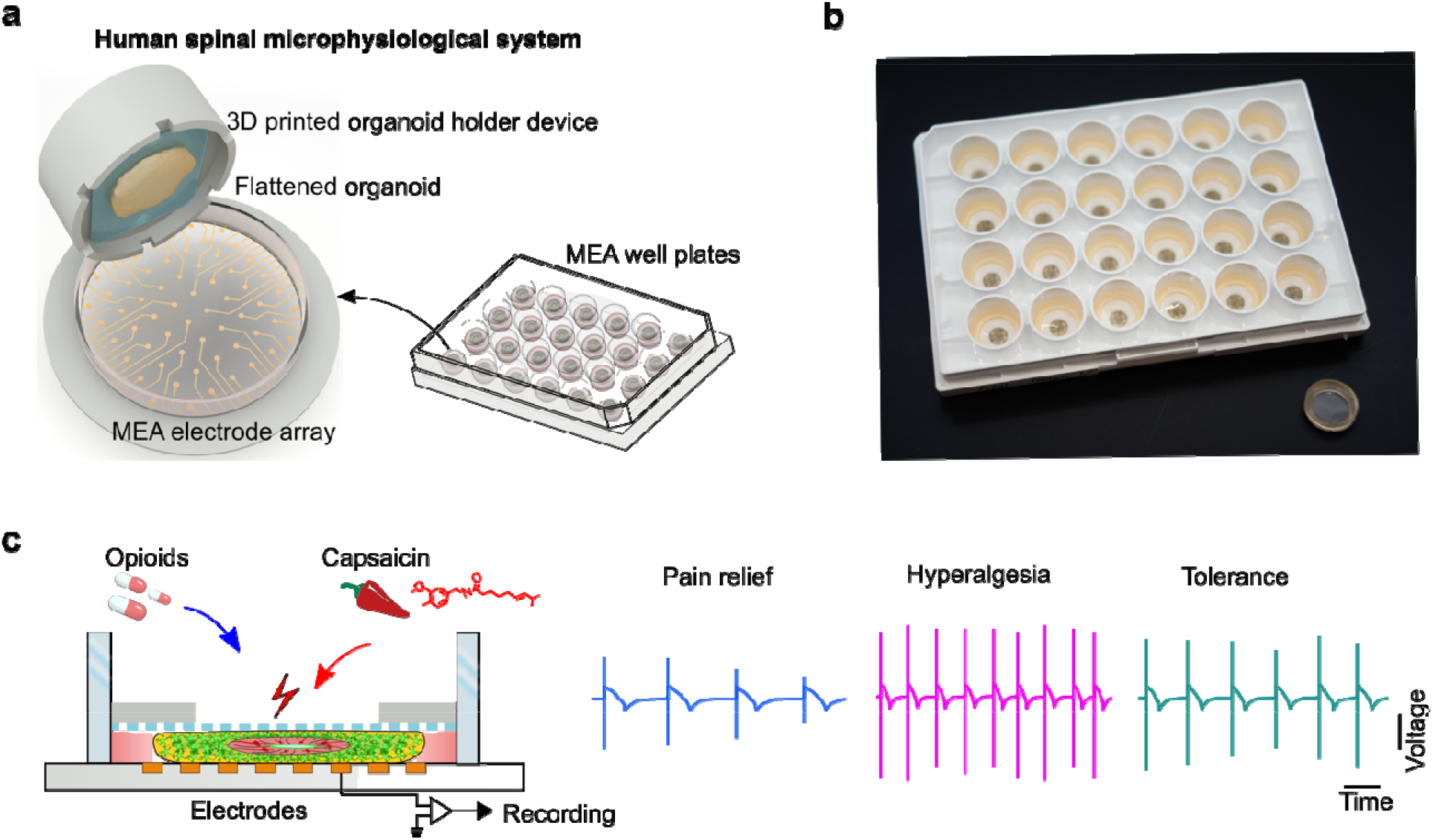
Engineering a human spinal microphysiological system to model opioid-induced tolerance and hyperalgesia. (**a**) Schematics of the human spinal microphysiological system that consists of a flattened human spinal cord organoid derived from human stem cells and a 3D printed organoid holder device. (**b**) A representative image of the human spinal microphysiological systems compatible with a well plate-based MEA system for plug-and-play neural activity measurement. The bottom right shows a single insert. (**c**) Schematic for modeling anti-nociception, opioid-induced hyperalgesia, and opioid tolerance using the human spinal microphysiological systems.

### Fabrication and characterization of flattened human spinal cord organoids

To fabricate flattened human spinal cord organoids, we developed a protocol that adapted previous dorsal spinal cord organoid protocols (**Figure S2**) and incorporated them into the organoid holder device. Briefly, we treated embryonic bodies (EBs) with retinoic acid and a GSK-3 inhibitor (CHIR-99021) for 4 days and then transferred them to a medium containing bone morphogen protein 4 (BMP4) and retinoic acid for directing dorsal spinal cord region specification for the next 6 days. Then, EBs/organoids were transferred to the organoid holder device, and organoids were treated with N-[N-(3,5-difluorophenacetyl)-l-alanyl]-S-phenyl glycine t-butyl ester (DAPT) for 8 days to promote neural differentiation. Finally, the organoids were kept in a medium supplied with cyclic adenosine monophosphate (cAMP) and ascorbic acid for further neural differentiation and maturation. We tracked organoid growth into the flattened shape within the confined space (height = 500 μm) between the device membrane and the well-plate bottom (**Figure 2a**). We further tested and compared hypoxia between conventional and flattened human spinal cord organoids (**Figure 2b**), and quantified the hypoxia within the two types of organoids (**Figure 2c**). We found a significantly reduced hypoxic core formation in the flattened organoids compared to the conventional organoids.

**Figure 2.**
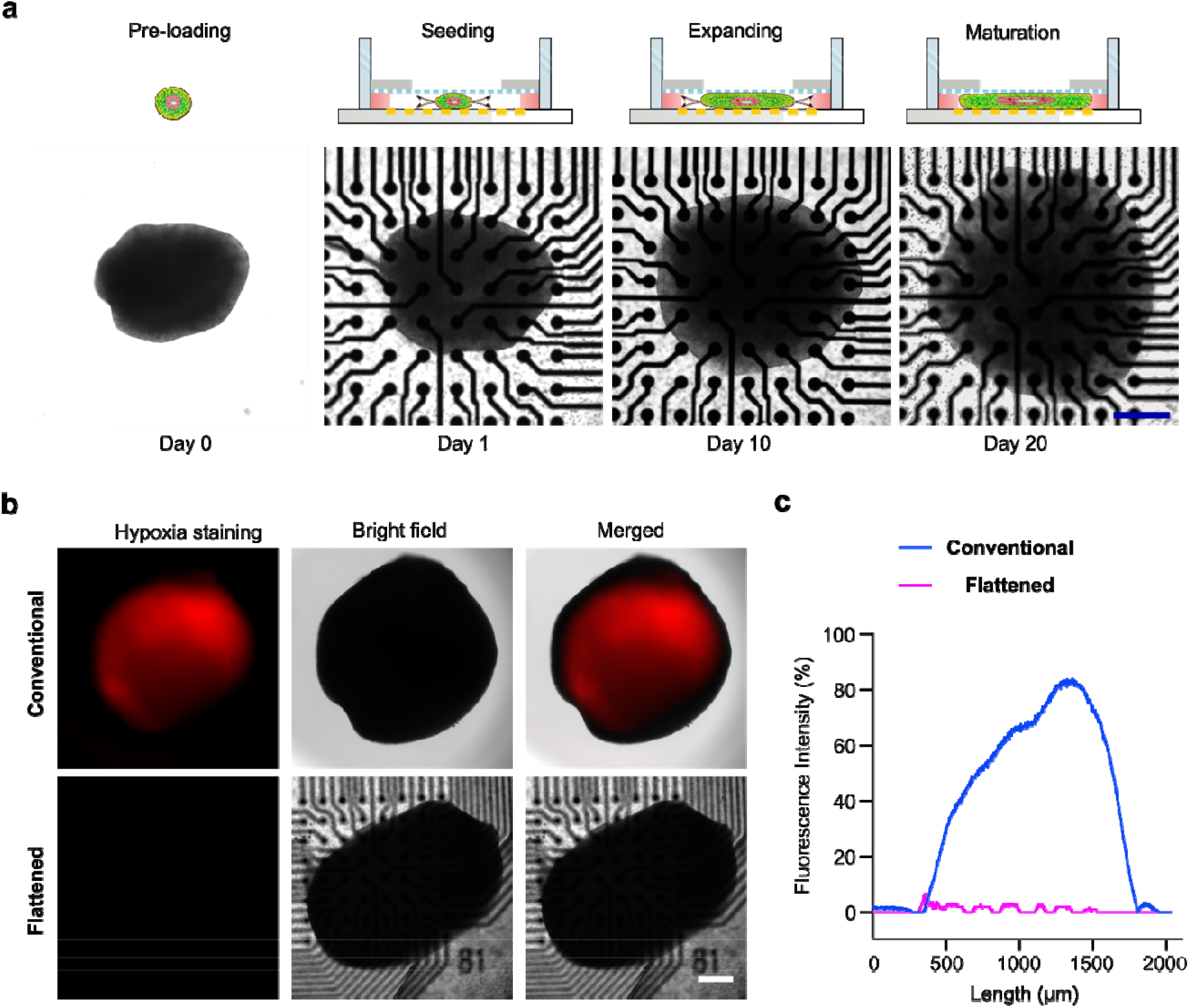
Fabrication and characterization of flattened human spinal cord organoids. (**a**) Schematics and images showing the fabrication process of the flattened human spinal cord organoids on-chip. (**b**) Hypoxia staining of the conventional and flattened human spinal cord organoids at 60 days. (**c**) Quantification of hypoxia within the conventional and flattened human spinal cord organoids at 60 days. Scale bar = 500μm.

### The flattened organoid design facilitates neural activity measurement and promotes neuron activity

The human spinal MPSs could be easily integrated into the MEA system for plug-and-play neural activity sensing. We showed that MPSs could be simply plugged into a neural activity measurement well of a 24 well MEA system (**Figure 3a**), and then tested right after plugging in. The conventional organoids showed a hypoxic center with no electrical activities, while the flattened organoids showed good neuron activities across the whole organoid area. While testing the neural activity of the flattened and conventional organoids, we obtained their representative raster plots (**Figure 3b**). In analyzing neural activity, we found the flattened organoids had significantly more active electrodes than the conventional organoids (**Figure 3c**), indicating that the flattened organoid design provides a better organoid-electrode contact for neural activity measurement. Moreover, we further quantified the mean firing rate of the organoids. We found that the mean firing rate of the flattened organoid (0.0658 ± 0.0234 Hz) was almost ten-fold higher than that of conventional organoids (0.0068 ± 0.0071 Hz, p=0.00082) (**Figure 3d**), implying that the human spinal MPSs have increased neural activity, perhaps due to the better perfusion afforded by the flattened organoid design. Furthermore, we characterized the neural activity of the human spinal MPSs during organoid development, with representative raster plots of human spinal MPSs obtained on days 10, 20, and 30 (**Figure 4a**). We also quantified the neural activity of the human spinal MPSs including burst frequency (**Figure 4b**), firing rate (**Figure 4c**), synchrony index (**Figure 4d**), and network burst frequency (**Figure 4e**). We found all the above neural activity parameters increased during organoid development, indicating that human spinal MPSs facilitated the development and maturation of neurons.

**Figure 3.**
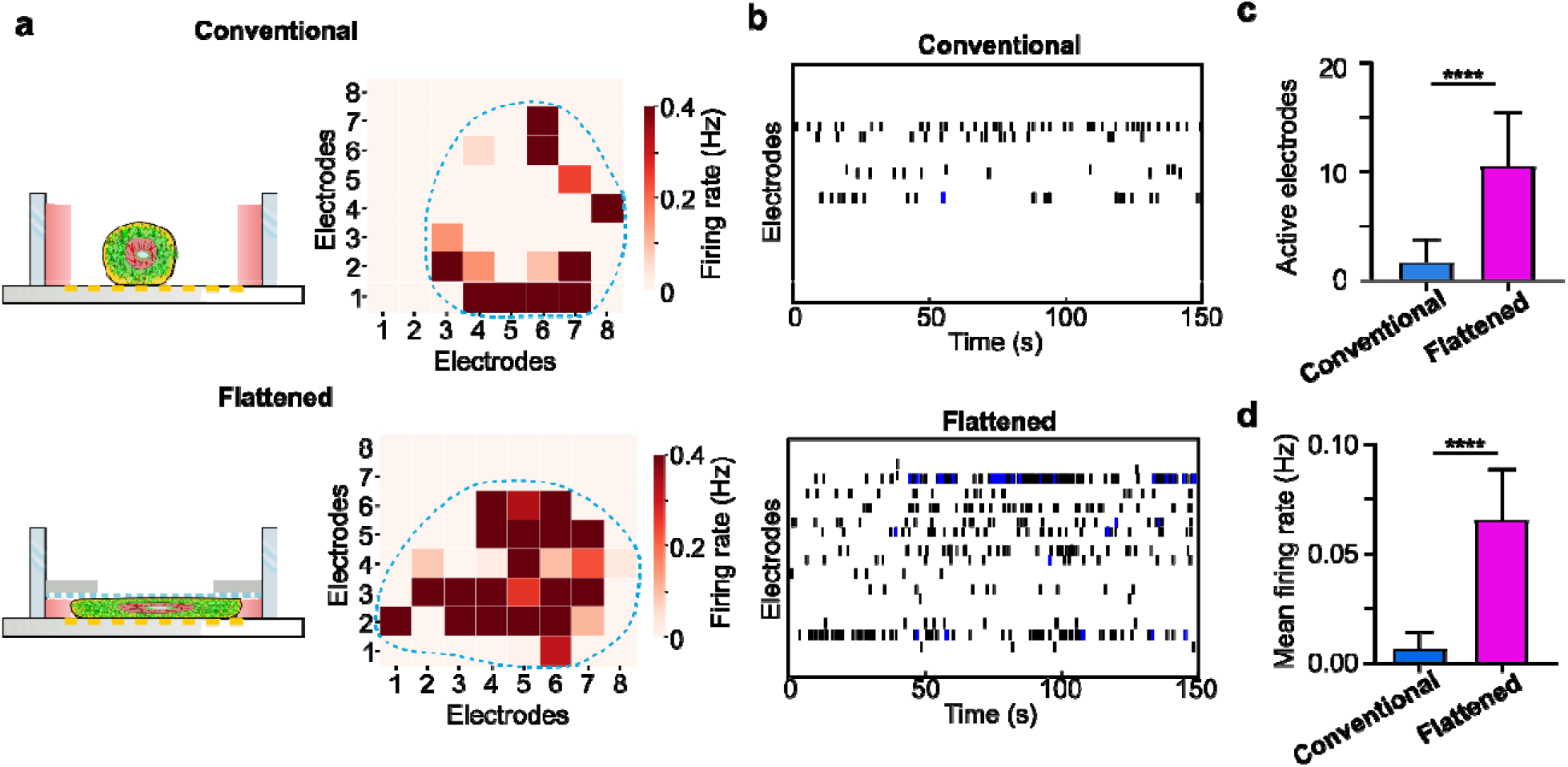
Neural activity measurements of conventional and flattened human spinal cord organoids. (**a**) Schematics and representative spike activity heat map of conventional and flattened human spinal cord organoids on the MEA plates. The blue dashed line labels the brain organoids area. (**b**) Representative raster plots of the conventional and flattened human spinal cord organoids. (**c**) Active electrode number of the conventional and flattened human spinal cord organoids during the neural activity recording. (**d**) Quantification of the mean firing rate of the conventional and flattened human spinal cord organoids (n=6). Scale bar = 500 μm.

**Figure 4.**
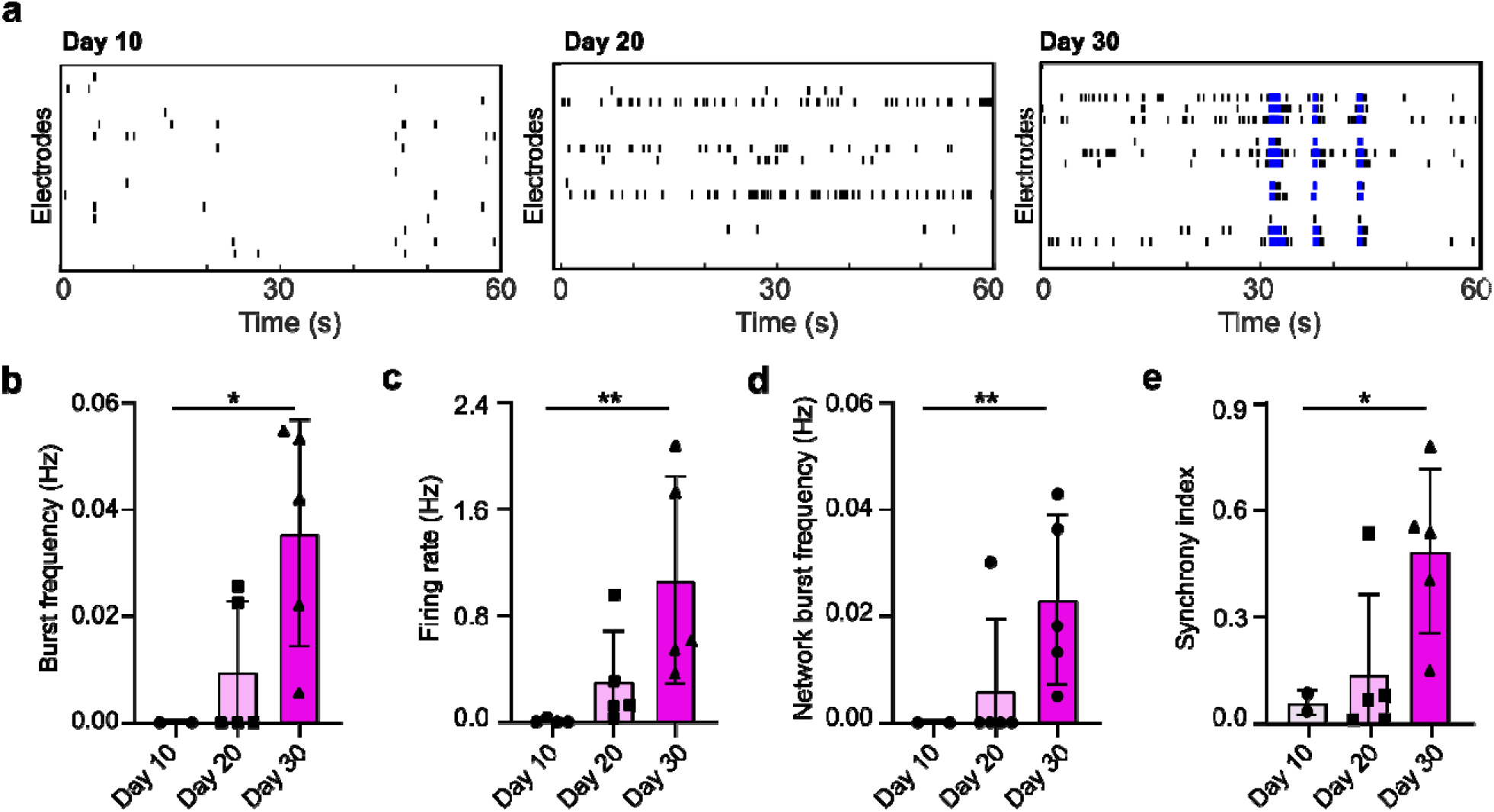
Functional maturation of human spinal microphysiological systems. (**a**) Representative raster plots of the human spinal microphysiological system at different developmental stages. The burst frequency (**b**), firing rate (**c**), synchrony index (**d**), and network burst frequency (**e**) of human spinal microphysiological systems at different ages in culture. (n=5) Scale bar = 500 μm.

### MPS response to short-term opioid administration

Opioids are widely used to relieve acute pain. To validate that the human spinal MPSs can be utilized to model the molecular mechanisms underlying pain relief during short-term opioid treatment, we first employed immunofluorescence staining to confirm the presence of neuronal populations that express μ-opioid receptors. The staining results showed that μ-opioid receptors were widely expressed in various types of neurons of human spinal MPSs, including excitatory, inhibitory, and sensory neurons (**Figure 5a**), similar to the findings from *in vivo* animal studies.^47-49^ To further validate the functional expression of μ-opioid receptors, we treated human spinal MPSs with increasing concentrations of the μ-opioid receptor agonist, DAMGO. DAMGO treatment inhibited organoid neural activity in a dose-dependent fashion (**Figure S3**). As the presence of μ-opioid receptors in neurons of human spinal MPSs was confirmed, we then tested the response of human spinal MPSs to pain modulators and opioid agonist treatments (**Figure 5b**). Capsaicin, a transient receptor potential cation channel subfamily V member 1 (TRPV1) agonist, was applied to MPSs to model nociception. Capsaicin stimulation increased the mean firing rate of organoids by 35.4%. Then we combined capsaicin and different concentrations of DAMGO to determine if opioids would decrease capsaicin-induced neuronal firing. The opioid treatment effectively reduced capsaicin responses and the mean firing rate of spinal cord organoids dropped below baseline even under a low dose of DAMGO (100 nM) (**Figure 5c, Video S1**). As the opioid concentration increased, the decrease of MPS firing rate became more apparent, and the organoid firing rate was minimal when the opioid concentration was greater than 1 μM. Thus, our MPSs could model capsaicin elicited pain activity and its treatment by opioids, recapitulating the in vivo situation.

**Figure 5.**
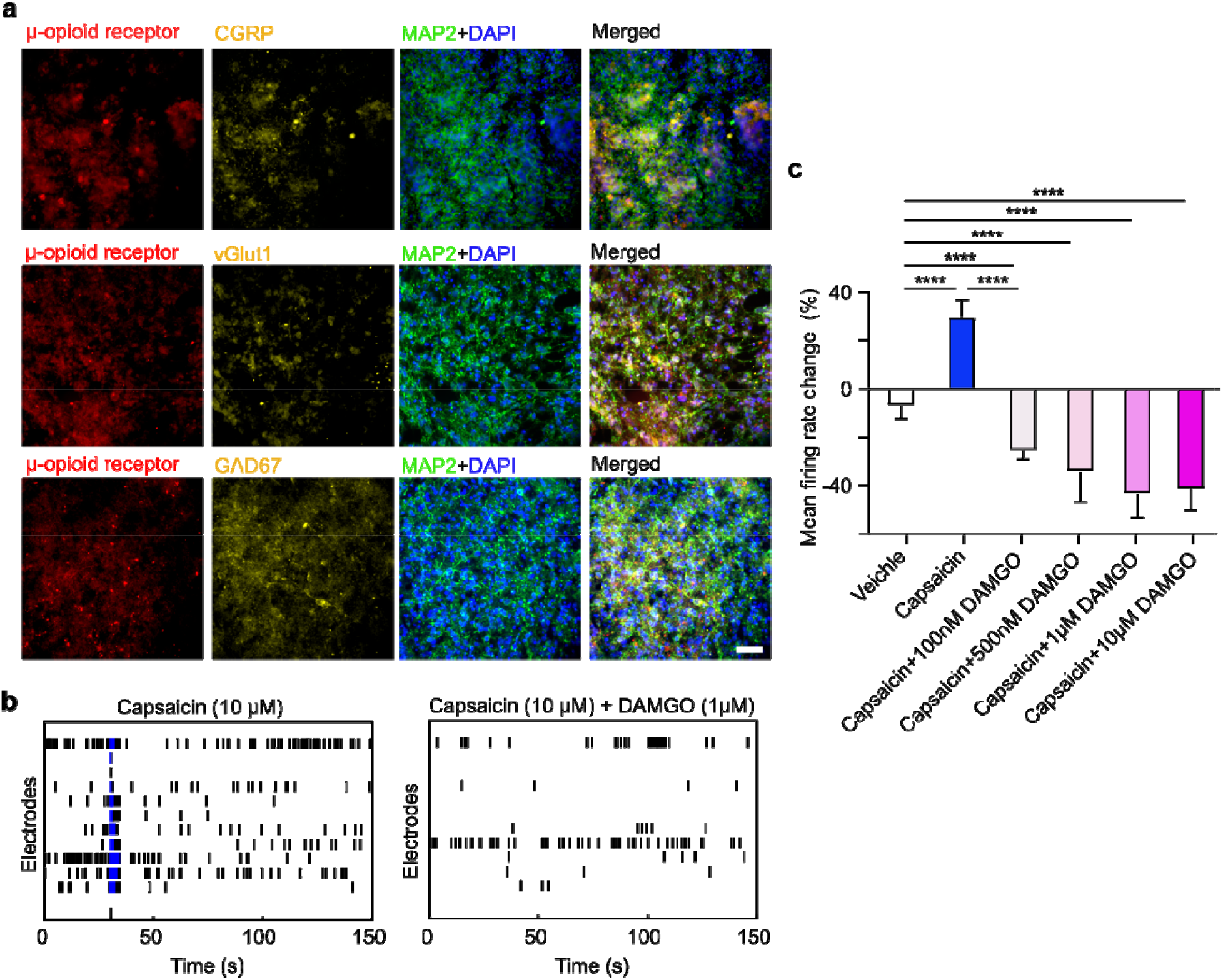
Model pain relief using human spinal microphysiological systems. (**a**) Immunofluorescence of human spinal microphysiological systems showing the expression of μ-opioid receptors on sensory neurons (CGRP), GABAergic (GAD67), and glutamate neurons (vGlut1). (**b**) Representative raster plots showing the neural activity signals of human spinal microphysiological systems treated with a noxious agent (capsaicin) only and capsaicin and opioid (DAMGO) together. (**c**) Percentage changes of mean firing rates after treatment with capsaicin only or by capsaicin with increasing concentrations of DAMGO (n=6). Scale bar = 50 μm.

### Chronic opioids induced hyperalgesia and tolerance

Extended opioid usage can induce a paradoxically increased sensitivity to pain (or opioid-induced tolerance) as well as tolerance to opioid-mediated pain relief. Both are major limitations when using opioids to treat chronic pain. To validate MPSs as a system for recapitulating opioid-induced hyperalgesia and tolerance observed following prolonged opioid administration, we continuously treated the human spinal MPSs with DAMGO (500 nM) for 8 days and tested their responses to pain modulators every other day (**Figure 6a, Video S2**). In the first 4 days, capsaicin increased the mean firing rate to an equivalent extent in the DAMGO-treated and the control groups. However, chronic DAMGO-treated organoids showed a significantly heightened firing rate increase on day 6 (70% ± 11% versus 51% ± 13%) and day 8 (87% ± 32% versus 39% ± 23%) of treatment. The results indicated a heightened sensitivity to capsaicin in human spinal MPSs during prolonged opioid administration. Thus, MSPs may be a good system for modeling opioid-induced tolerance. We also tested whether organoids treated together with DAMGO and naloxone, a μ-opioid receptor antagonist, could reduce neuronal activity in the model of opioid-induced hyperalgesia. The DAMGO+naloxone group showed no significant difference in capsaicin-induced mean firing change over eight days, indicating a requirement of mu-opioid receptors in this model of hyperalgesia. We further measured the responses of human spinal MPS to capsaicin and DAMGO (with different doses) to test opioid tolerance after extended opioid administration (**Figure 6b**). A low dose of DAMGO (100 nM) was able to reduce the firing rate below the baseline and effectively relieve the neuronal firing induced by capsaicin in the control and DAMGO+Naloxone group, while organoids chronically treated with DAMGO needed higher concentrations of DAMGO to return neuronal activity to baseline. The results from the human spinal MPSs are similar to those findings on spinal cord slices from animals with both opioid-induced hyperalgesia and tolerance. We further validated human spinal MPSs as able to recapitulate the key receptor and pathological changes with prolonged opioid administration via immunofluorescence staining of the three groups (**Figure 6c**). We found that μ-opioid receptor expression was downregulated after prolonged DAMGO treatment.

**Figure 6.**
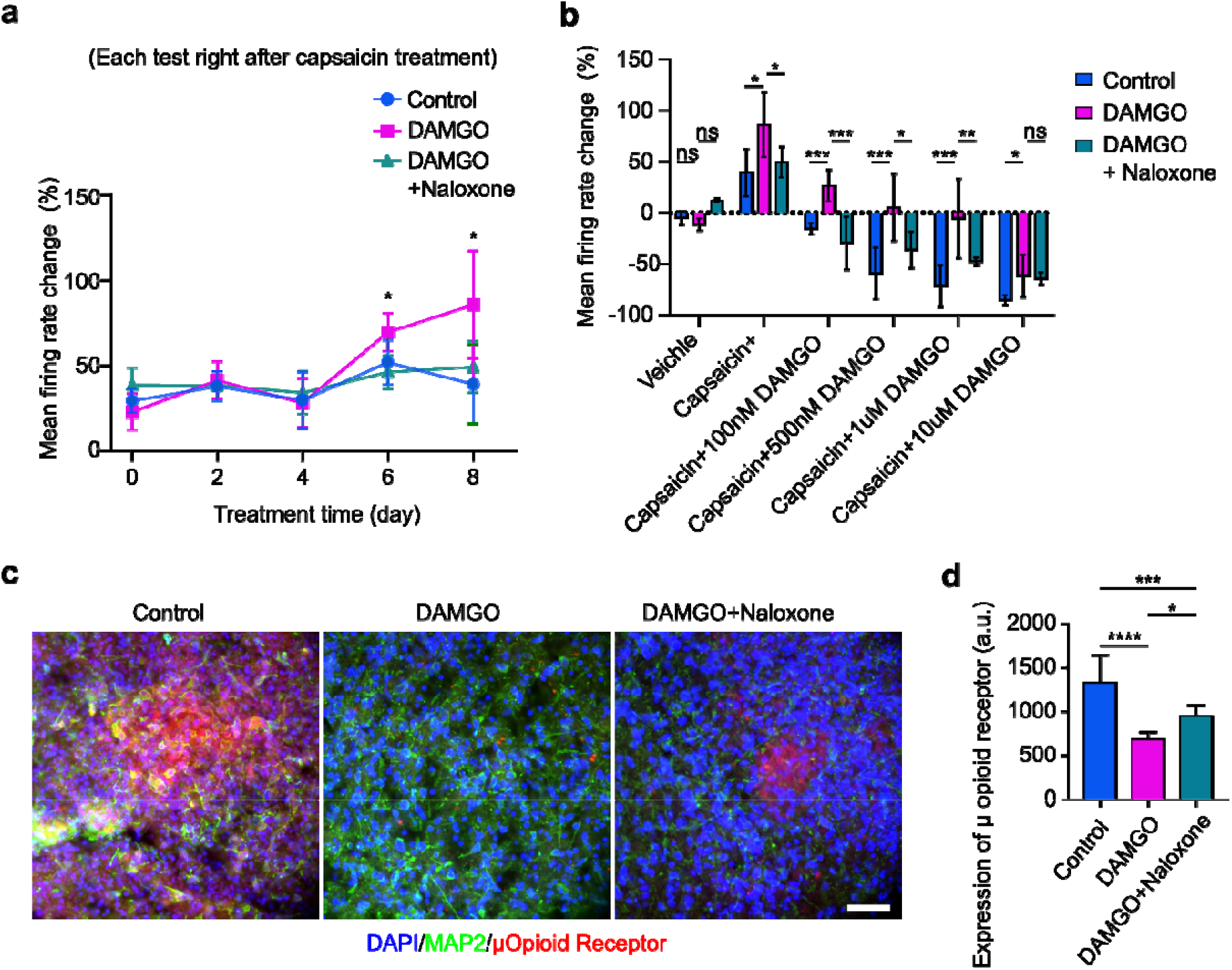
Modeling opioid-induced hyperalgesia and µ-opioid receptor down-regulation using human spinal microphysiological systems. (**a**) Percentage change of mean firing rates for the three groups including vehicle treatment only (Control), opioid treatment only (500 nM DAMGO), and the combination of opioid and naloxone treatment (500 nM DAMGO + 1 μM naloxone) over 8 days. (**b**) After the 8-day treatment, the percentage changes of mean firing rates from baseline by treating with a noxious agent (capsaicin), and capsaicin together with different doses of DAMGO. (**c**) Immunofluorescence showing the expression of μ-opioid receptors within human spinal microphysiological systems for the three groups after the 8-day treatment. (**d**) Quantification of μ-opioid receptor expression within the three groups as measured by fluorescent intensity (n = 6). Scale bar = 50 μm.

## 3. Conclusion

In summary, we developed novel human spinal microphysiological systems (MPSs) integrated with neural activity sensing for modeling human opioid-induced analgesia, tolerance and hyperalgesia. The MPSs have several unique advantages: (1) the innovative flattened organoid design of MPSs enhances nutrient and oxygen entry to avoid typical organoid hypoxia/necrosis and to support neuron maturation/activity; (2) the flattened organoid design of MPSs also provides better contact between the organoid and the electrodes from a multi-electrodes array (MEA), improving the stability and reproducibility of neural activity measurement; (3) the MPSs in the 3D printed organoid holder devices are compatible with commonly-used well-plates or MEA systems, highlighting their potential for broad applications in basic and translational pain research. Using this model, we not only demonstrated that acute opioid exposure could rapidly reduce neural activity evoked by capsaicin, but also showed that prolonged opioid exposure downregulates μ-opioid-receptor levels as well as glutamate transporter levels and produces neurophysiological and neurochemical features associated with both tolerance and opioid-induced hyperalgesia. We believe the human spinal MPSs could be broadly applied to studies of pain etiology and its treatment (e.g. assessments of biased agonism, receptor internalization, etc.). Successful implementation of human spinal microphysiolgocial systems will facilitate drug discovery efforts aimed at identifying and validating new pain therapeutics while enhancing prospects for successful clinical translation.

## 4. Experimental Section

### Fabrication of 3D printed organoid holder devices

The organoid holder devices were designed employing AutoCAD software and fabricated using our well-developed 3D printing protocol^50, 51^ through a stereolithography 3D printer (Form 3B, Formlabs) and FormLabs Clear Resin V4. The detailed device design is shown in **Figure S2a**.

### Human stem cell culture

We obtained human embryonic stem cells (WA09) from WiCell institute and followed the guidelines of both WiCell Institute and Indiana University when handling these cells. Matrigel (Corning) coated 6 well plates were used to culture WA09 cells with mTESR plus medium (Stemcell Technologies) in an incubator at 37 degrees Celsius and 5% CO_2_. The medium was changed every other day. ReLeSR (Stemcell Technologies) was used to passage WA09 cells weekly.

### Fabrication of flattened human spinal cord organoids

Similarly, to our early protocols,^52, 53^ embryonic bodies (EBs) were fabricated by aggregation of ∼9,000 WA09 cells each well using a 96-well U bottom microplate (Corning). EBs were cultured in 100 μL EB formation medium (Stemcell Technologies) supplemented with 10 µM Y-27632 (SelleckChem). After one day of EB formation, the EBs were transferred to the spinal cord medium I (ScM I) supplied with 10 nM retinoic acid (Sigma Aldrich) and 3 μM CHIR-99021 (Stemcell Technologies). After 4-day culture in SCM I, the spheroids were then transferred to spinal cord medium II (ScM II) with 10 nM retinoic acid and 5 ng/mL recombinant human BMP4 (Peprotech) for the next 6 days. On day 10, we embedded the spinal cord organoids into Matrigel (corning). The organoids were switched to spinal cord medium III (ScM III), supplemented with 10 μM DAPT, and cultured for 8 more days. During this period, spinal cord organoids were transferred to our microfluidic devices integrated with 24 well ultra-low attachment plates (Corning) or 24 well MEA systems for the generation of flattened organoids. For traditional spherical organoids, the organoids were cultured in 6 well ultra-low attachment plates (Corning) shaking at 60RPM with an orbital shaker (OrbiShaker JR, Benchmark Scientific). On Day 18, the organoids were finally transferred to spinal cord medium IV (ScM IV), with 1 μM cAMP (Sigma Aldrich) and 20 μg/mL ascorbic acid (Sigma Aldrich) for subsequent continuous culture. Medium change was performed every other day during this process except when specified. Detailed medium composition is listed in **Supplementary Table S1**.

### Neural activity measurement of human spinal cord organoids

On Day 10, the human spinal cord organoids were transferred into devices. After having settled for 10 days, human spinal cord organoids were attached to wells on the MEA plates and prepared for electrical measurements. Recording of human spinal cord organoids’ electrical activity was performed by using the Axion Maestro Edge (Axion inc). The spinal cord organoids were kept at 37°C maintained by an internal heater, and infused with 5% CO_2_. To determine the mean firing rate of a spinal cord organoid, we repeated 3-minute MEA recordings 5 times after the treatment administration. The final mean firing rate of the organoid was calculated as the average mean firing rate of the 3 most consistent recordings among the 5 recordings. To reduce the variation brought by adding substance, the medium was also refreshed before measuring the mean firing rate baseline. The substance was first diluted into SCM IV at 2x of the desired concentration, then half of the medium was replaced with the 2x concentrated solution to reach the desired concentration.

### Cryo-section of organoids

After treating with different drugs as mentioned, the culture samples were washed three times in 1x Phosphate-Buffered Saline (PBS, Gibco) and then fixed in 4% paraformaldehyde in 1x PBS (Thermo Scientific) overnight at 4 °C. The samples were rinsed three times with 1x PBS and then transferred into a 30% sucrose (w/v) solution overnight at 4 ºC for cryoprotection. The organoids were then incubated with 7.5% gelatin (w/v) and 10% sucrose (w/v) in PBS solution at 37 ºC for 1 hour. Finally, the samples were snap-frozen onto a cryomold (Sakura Finetek) in a dry ice/ethanol slurry. The frozen samples were then sectioned at 30 µm thickness on a cryostat (Leica).

### Immunofluorescence staining

We characterized the human spinal cord organoids using protocols previously developed in the lab^54-57^. The cryosectioned samples plated onto glass slides were washed three times with 1x PBS. Then, for antigen retrieval, the slides were treated with 2N hydrochloric acid (HCl) for 15 minutes. After HCl treatment, the samples were further washed twice using 1XPBS and treated with a blocking solution (0.3% Triton-X100, 5% normal goat serum in 1XPBS) for 1 hour. Following blocking, the samples were immersed in a blocking solution containing primary antibodies at 4 ºC overnight. We then washed the sections 3 times with 1x PBS, and incubated the samples with a blocking solution containing secondary antibodies for 1 hour at room temperature. Finally, the samples were washed with 1x PBS another 3 times and coverslipped with gold anti-fade mounting medium (Invitrogen). Detailed antibody information and dilution factors can be found in **Supplementary Table S2**.

### qPCR analysis

Mature organoids were lysed for RNA extraction using RNeasy Mini Kit (Qiagen). RNA was first reversed and transcribed into complementary DNA (cDNA) using a cDNA synthesis kit (Quantabio). The cDNA was then analyzed by quantitative PCR using the SYBR green kit (Applied Biosystems). Gene expression fold change was analyzed by normalizing against the housekeeping gene GAPDH.

### Statistical analysis

Statistical analysis was carried out using GraphPad Prism 8. Two sample groups were compared using the paired Students’ t-test. Statistical significance was denoted as: *p<0.05, **p<0.01, ***p<0.005. ****p<0.001.

## 5. Acknowledgment

The project was supported by the departmental start-up funds of Indiana University Bloomington, and in part by the NSF grant (CCF-1909509) and NIH awards (DP2AI160242, DA056242, and DA047858).

